# Very severe anemia and one year mortality outcome after hospitalization in Tanzanian children: A prospective cohort study

**DOI:** 10.1101/581462

**Authors:** Neema Chami, Duncan K. Hau, Tulla S. Masoza, Luke R. Smart, Neema M. Kayange, Adolfine Hokororo, Emmanuela E. Ambrose, Peter P. Moschovis, Matthew O. Wiens, Robert N. Peck

**Affiliations:** Department of Pediatrics, Catholic University of Health and Allied Sciences, Mwanza, Tanzania; Department of Pediatrics, Weill Cornell Medical College, New York, NY, USA; Division of Hematology/Oncology, Department of Pediatrics, Cincinnati Children’s Hospital Medical Center, Cincinnati, OH, USA; Divisions of Pediatric Global Health and Pulmonary Medicine, Department of Pediatrics, Massachusetts General Hospital, Boston, MA, USA; Faculty of Medicine, Mbarara University of Science and Technology, Mbarara, Uganda; Center for International Child Health, BC Children’s Hospital & University of British Columbia, Vancouver, Canada; Center for Global Health, Weill Cornell Medical College, New York, NY, USA

## Abstract

**Background:** Africa has the highest rates of child mortality. Little is known about outcomes after hospitalization for children with very severe anemia.

**Objective:** To determine one year mortality and predictors of mortality in Tanzanian children hospitalized with very severe anemia.

**Methods:** We conducted a prospective cohort study enrolling children 2-12 years hospitalized from August 2014 to November 2014 at two public hospitals in northwestern Tanzania. Children were screened for anemia and followed until 12 months after discharge. The primary outcome measured was mortality. Predictors of mortality were determined using Cox regression analysis.

**Results:** Of the 505 children, 90 (17.8%) had very severe anemia and 415 (82.1%) did not. Mortality was higher for children with very severe anemia compared to children without over a one year period from admission, 27/90 (30.0%) vs. 59/415 (14.2%) respectively (Hazard Ratio (HR) 2.42, 95% Cl 1.53–3.83). In-hospital mortality was 11/90 (12.2%) and post-hospital mortality was 16/79 (20.2%) for children with very severe anemia. The strongest predictors of mortality were age (HR 1.01, 95% Cl 1.00–1.03) and decreased urine output (HR 4.30, 95% Cl 1.04 – 17.7).

**Conclusions:** Children up to 12 years of age with very severe anemia have nearly a 30% chance of mortality following admission over a one year period, with over 50% of mortality occurring after discharge. Post-hospital interventions are urgently needed to reduce mortality in children with very severe anemia, and should include older children.

## Introduction

Very severe anemia (hemoglobin concentration less than 5.0 g/dL) is a major cause of morbidity and mortality among children in Africa. Of hospitalized children, 12 to 29% have very severe anemia, and the in-hospital mortality of these children range between 4 to 17% [1–9]. Little is known about the long-term outcomes for children with very severe anemia after hospitalization [10]. The few data that have been published have been limited to children under five years of age [9]. Studies are lacking regarding long-term outcomes after hospitalization that include older children with very severe anemia.

Therefore, we conducted a prospective cohort study of Tanzanian children up to 12 years of age hospitalized with very severe anemia and followed until one-year after hospital discharge. Our study objectives were: 1) to determine the prevalence of very severe anemia for hospitalized children, 2) to compare mortality in children with very severe anemia to children without very severe anemia up to one year post-hospitalization, and 3) to identify predictors of mortality for children with very severe anemia.

## Methods

### Study site

This study is a secondary analysis of a prospective cohort study where we consecutively screened and enrolled children hospitalized on the pediatric wards of two public hospitals in northwestern Tanzania and followed for one-year post hospitalization [11]. Bugando Medical Center (BMC) and Sekou-Toure Regional Referral Hospital (STH) are in the city of Mwanza, the second largest city in Tanzania and the capital of the Mwanza region. BMC is a public tertiary hospital that serves as the zonal referral hospital for northwestern Tanzania with a catchment area of approximately 16 million people. BMC has 1,000 inpatient beds and approximately 3,500 pediatric hospitalizations per year. STH is a public regional hospital for Mwanza region with a population of approximately 3 million people. STH has 320 inpatient beds and approximately 2,000 pediatric hospitalizations per year.

### Inclusion and exclusion criteria

Children 2-12 years of age hospitalized in the medical ward of BMC or STH were eligible for enrollment in the study. The parent or guardian of a potential participant was provided with information regarding the study within 12 hours of admission. Children were enrolled only after obtaining informed consent from a parent or guardian by a study member. Study participants with multiple hospitalizations to BMC or STH during the study period were only enrolled during their first hospitalization. Children who were referral cases from another hospital were excluded from the study.

### Study procedures

On the day of enrollment, a modified version of the World Health Organization (WHO) STEPS questionnaire was administered in Swahili by a Tanzanian study investigator to the parent or guardian [12]. The WHO STEPS questionnaire includes questions regarding socioeconomic status, medical history, prior testing, diagnosis and treatment for diseases as well as standard protocols for physical examination. After completing the questionnaire, the study investigator conducted a standardized physical examination including the measurement of vital signs, weight and height. An axillary temperature was taken. Weight was measured to the nearest 0.1 kg using a digital scale (DETECTO, USA), which was adjusted to zero before each measurement. If the child was unable to stand, her weight was taken on a hanging scale. Height was measured to the nearest 0.1 cm using a stadiometer. If the child was unable to stand, her height was measured while lying down.

All children underwent measurements of hemoglobin, glucose, creatinine, and urine dipstick testing as standard procedures of hospitalization at BMC and STH. Hemoglobin levels were measured by Hemo Control Hemoglobin Analyzer (EKF Diagnostics, Germany). Glucose levels were measured by Ascensia Glucometer (Bayer Healthcare, Germany). Serum creatinine levels were measured using Cobas Integra 400 Plus Analyzer (Roche Diagnostic Limited, Switzerland). An estimated glomerular filtration rate (eGFR) was calculated using the bedside Schwartz equation as recommended by international guidelines [13]. A urine dipstick was used to test for proteinuria and hematuria (Multistix 10SG, Siemens, USA). At the time of hospitalization, by national policy, all children were offered testing for human immunodeficiency virus (HIV) according to the Tanzania national guidelines [14].

### Study definitions

Severity of anemia was defined according to WHO reference standard for anemia [15]. Study participants with a hemoglobin concentration less than 5.0 g/dL were classified as very severe anemia.

### Clinical procedures and diagnoses

Disease management was conducted by the hospital clinicians in accordance with hospital and Tanzanian management protocols. Per standard hospital policy, children with very severe anemia received an urgent blood transfusion (whole blood 20 milliliters per kilogram body weight).

BMC and STH have used a standard list of recommended pediatric diagnoses adapted from the WHO’s International Classification of Disease version 10 (ICD-10) [16]. For a child with multiple diagnoses (e.g., severe malnutrition and diarrheal disease), a single diagnosis was recorded for this study that was based on the primary diagnosis recorded by the clinicians caring for the child. Caretakers were given standard discharge instructions and told to follow-up in clinic within 2 weeks of discharge or sooner if necessary.

### Follow-up of study participants

A total of three mobile phone numbers were obtained from all participants’ caretakers at the time of discharge. This included one number for the study participant’s parent or guardian, and two additional numbers for relatives or close friends. Follow-up phone calls were made at 3, 6 and 12 months post-discharge. If the child had died, the date of death was also determined.

### Study outcome

The primary study outcome was mortality. Mortality was classified as in-hospital if it occurred during the index hospitalization and post-hospital if it occurred in the year that followed the index hospitalization.

### Data analysis

Data were entered into Microsoft Excel (Microsoft, Redmond, Washington, USA) and analyzed using Stata version 14 (College Station, Texas, USA). Categorical variables were described as proportions (percentages), and continuous variables were described as means (standard deviations). For all cross-sectional analyses, a chi-squared test was used for comparing categorical variables and a Wilcoxon rank sum test was used for continuous variables. Cox regression models were used for all survival analyses to compare outcomes between study groups and to determine predictors of mortality. All variables with *p* < 0.05 in the univariable model along with age and sex were entered into the multivariable model. Kaplan-Meier survival curves were used to display incident mortality. A log-rank test was used to determine if mortality incidence differed by severity of anemia. Study participants lost to follow-up were censored at the last contact date. All available data were included in all calculations. No variable was missing for more than 14 participants. A two-sided *p*-value of < 0.05 was regarded as statistically significant in all analyses.

### Ethical consideration

Permission to conduct the study was obtained from the research committees of Bugando Medical Center, Sekou-Toure Regional Referral Hospital, Weill Cornell Medical College, and the National Institute for Medical Research in Tanzania. Participants were enrolled only after obtaining informed consent from one of their parents or guardian. Parents also consented to receive phone calls at either their own mobile phone number or the mobile phone numbers they provided for relatives or friends. They agreed that if they were not available to receive the phone call, relatives or friends could provide information about the vital status of the child. All study procedures were in accordance with the ethical standards of the responsible committee on human experimentation (institutional and national) and with the Helsinki Declaration of 1975, as revised in 2000.

## Results

### Study enrollment

From 1^st^ August 2014 to 30^th^ November 2014, 537 children were hospitalized in the pediatric wards of BMC and STH. Of the 537 children hospitalized, 15 died before enrollment occurred, 8 were excluded for being referral cases from another hospital, and 8 declined participation. The remaining 506 children (94.2%) were enrolled, with 461/506 (91.1%) at BMC and 45/506 (8.9%) at STH. One participant of the 506 children did not have a hemoglobin level measured.

Of the 505 children, 90 (17.8%) had very severe anemia and 415 (82.1%) did not. The in-hospital mortality rates were 11/90 (12.2%) for children with very severe anemia and 28/415 (6.7%) for children without very severe anemia. Of the 466 children who were discharged from the hospital, mobile phone contact was made with 458/466 (98.3%) children’s parents (or designated proxies) at 3 months, 409/466 (87.8%) at 6 months, and 372/466 (79.8%) at 12 months. Twenty-two of 79 (27.8%) children with very severe anemia and 72/387 (18.6%) children without very severe anemia were lost to follow-up at 12 months.

### Baseline characteristics

The baseline characteristics of the two study groups are described in Table 1. Children with very severe anemia were found to be older (59.1 vs 53.6 months, *p* = 0.027). Other notable differences were significantly less reports of fever, more reports of taking herbal medication, lower diastolic blood pressure, lower Glasgow coma scores, more children with pallor, higher eGFR, more children with hematuria, more children with sickle cell disease, and fewer children with diarrheal, respiratory, urinary tract infection or neurologic diseases. The mean hemoglobin level for children with very severe anemia was 3.6 g/dL compared to 8.8 g/dL for children without very severe anemia.

**Table 1.**
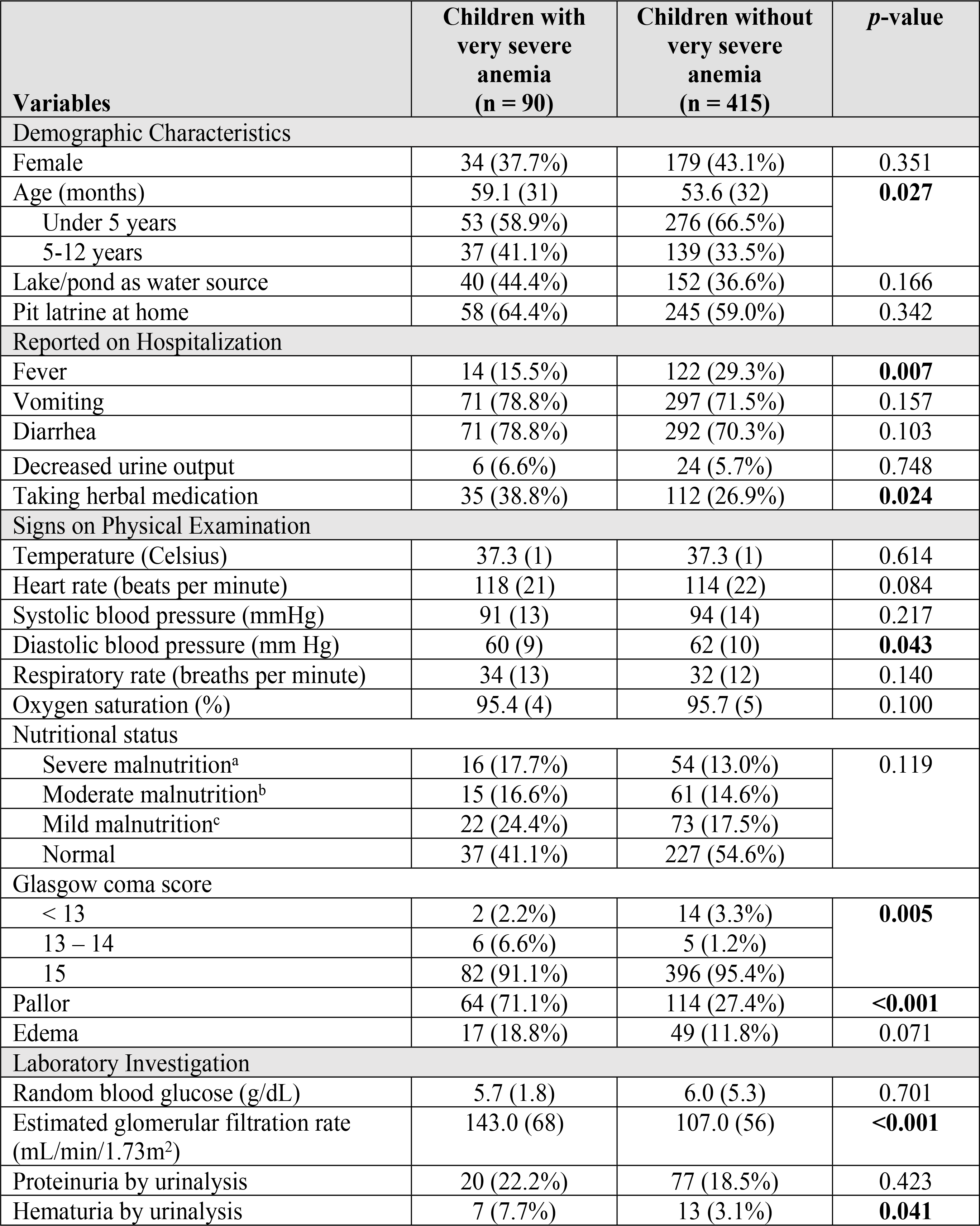
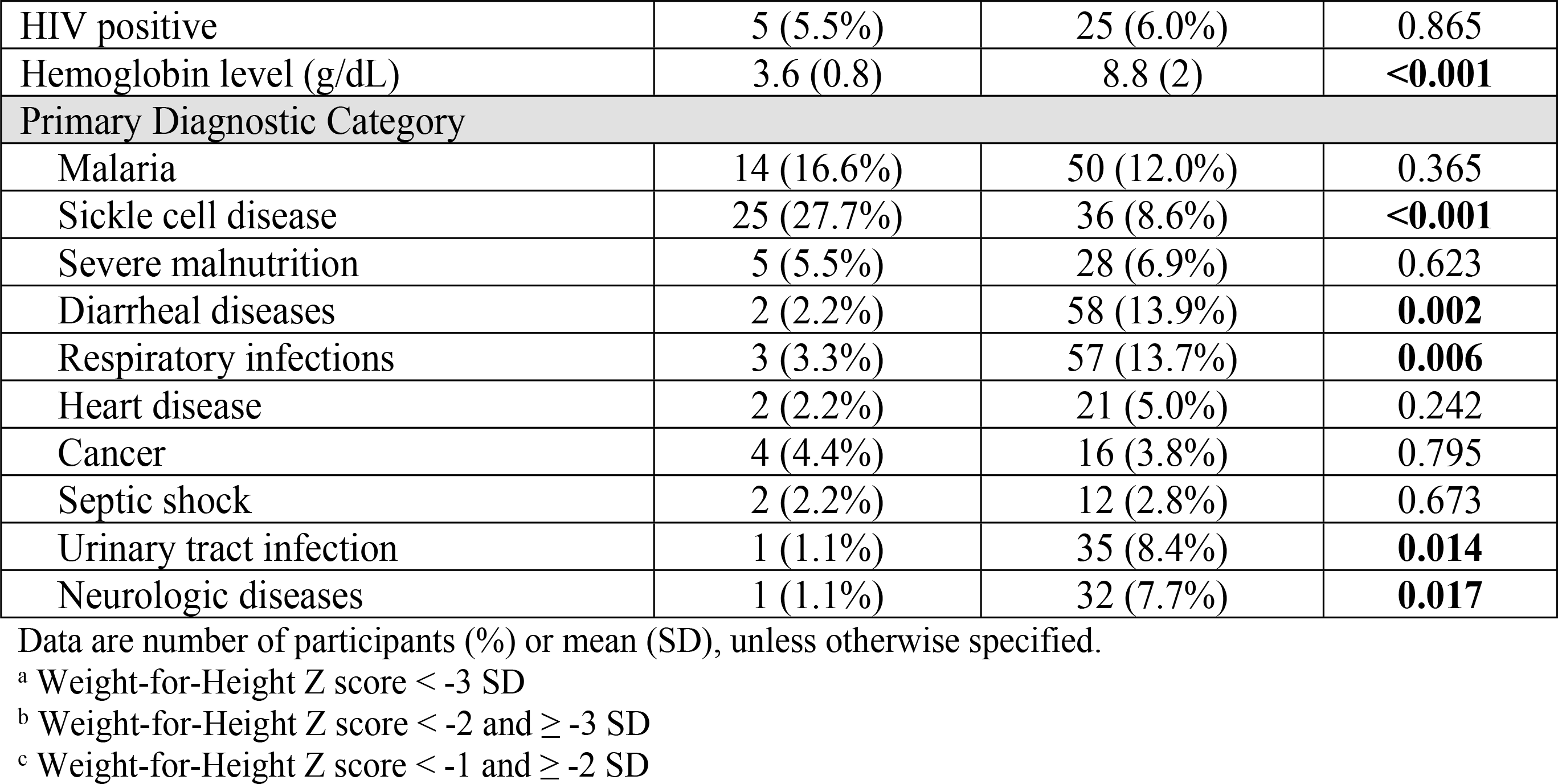
Baseline characteristics of children with very severe anemia and without very severe anemia.

### Prevalence of very severe anemia

Among all participants, 90/505 (17.8%) were found to have a hemoglobin level of less than 5.0 g/dL upon admission (Table 2). A total of 203/505 (40.2%) children had either severe or very severe anemia. The total number of children that had any level of anemia was 442/505 (87.5%).

**Table 2.** Severity of anemia and overall mortality at 12 months for all study participants.

|  | Total<br>(N = 505) | Dead<br>(N = 86) | Alive<br>(N = 419) | HR<br>[95% CI] | p-value |  |
| --- | --- | --- | --- | --- | --- | --- |
| Hemoglobin level (g/dL) | 7.9<br>(2.7) | 6.5<br>(2.6) | 8.1<br>(2.6) | 0.82<br>[0.75 – 0.88] | <0.001 |  |
| Anemia severity<sup>a</sup> |  |  |  |  |  |  |
| Very severe anemia | 90<br>(17.8%) | 27<br>(30.0%) | 63<br>(70.0%) | 4.33<br>[1.66 – 11.26] | 0.003 | <0.001<sup>b</sup> |
| Severe anemia | 113<br>(22.4%) | 24<br>(21.2%) | 89<br>(78.8%) | 2.77<br>[1.06 – 7.28] | 0.038 |  |
| Moderate anemia | 187<br>(37.0%) | 24<br>(12.8%) | 163<br>(87.2%) | 1.55<br>[0.59 – 4.07] | 0.370 |  |
| Mild anemia | 52<br>(10.3%) | 6<br>(11.5%) | 46<br>(88.5%) | 1.50<br>[0.45 – 4.92] | 0.502 |  |
| No anemia | 63<br>(12.5%) | 5<br>(7.9%) | 58<br>(92.1%) | Reference |  |  |

Data are number of participants (%) or mean (SD)
<sup>a</sup> Defined according to WHO reference standard for anemia
<sup>b</sup> p trend

### Mortality

Overall one-year mortality was significantly higher for children with very severe anemia compared to children without very severe anemia, 27/90 (30.0%) vs. 59/415 (14.2%) respectively (Hazard Ratio [HR] 2.42, 95% Cl 1.53 – 3.83, *p* < 0.001). In-hospital mortality was higher for children with very severe anemia compared to those without very severe anemia, 11/90 (12.2%) vs. 28/415 (6.7%) respectively (HR 1.89, 95% Cl 0.94 – 3.79, *p* = 0.073). Post-hospital mortality was significantly higher for children with very severe anemia compared to those without very severe anemia, 16/79 (20.2%) vs. 31/387 (8.0%) respectively (HR 2.98, 95% Cl 1.62 – 5.45, *p* < 0.001). For post-hospital mortality, the median time-point for death among children with very severe anemia was 57 days after discharge, compared to 190 days after discharge for those without very severe anemia.

Children with very severe anemia had a 4.3-fold increase risk of mortality compared to children without any anemia (HR 4.33, 95% Cl 1.66 – 11.26, *p* = 0.003) (Table 2). As the severity of anemia increased, the hazards of mortality increased (Fig 1). For every 1 g/dL decrease in hemoglobin level, the risk of death increased by 18% for all study participants (HR 0.82, 95% Cl 0.75 – 0.88, p < 0.001) (Table 2).

**Fig 1.**
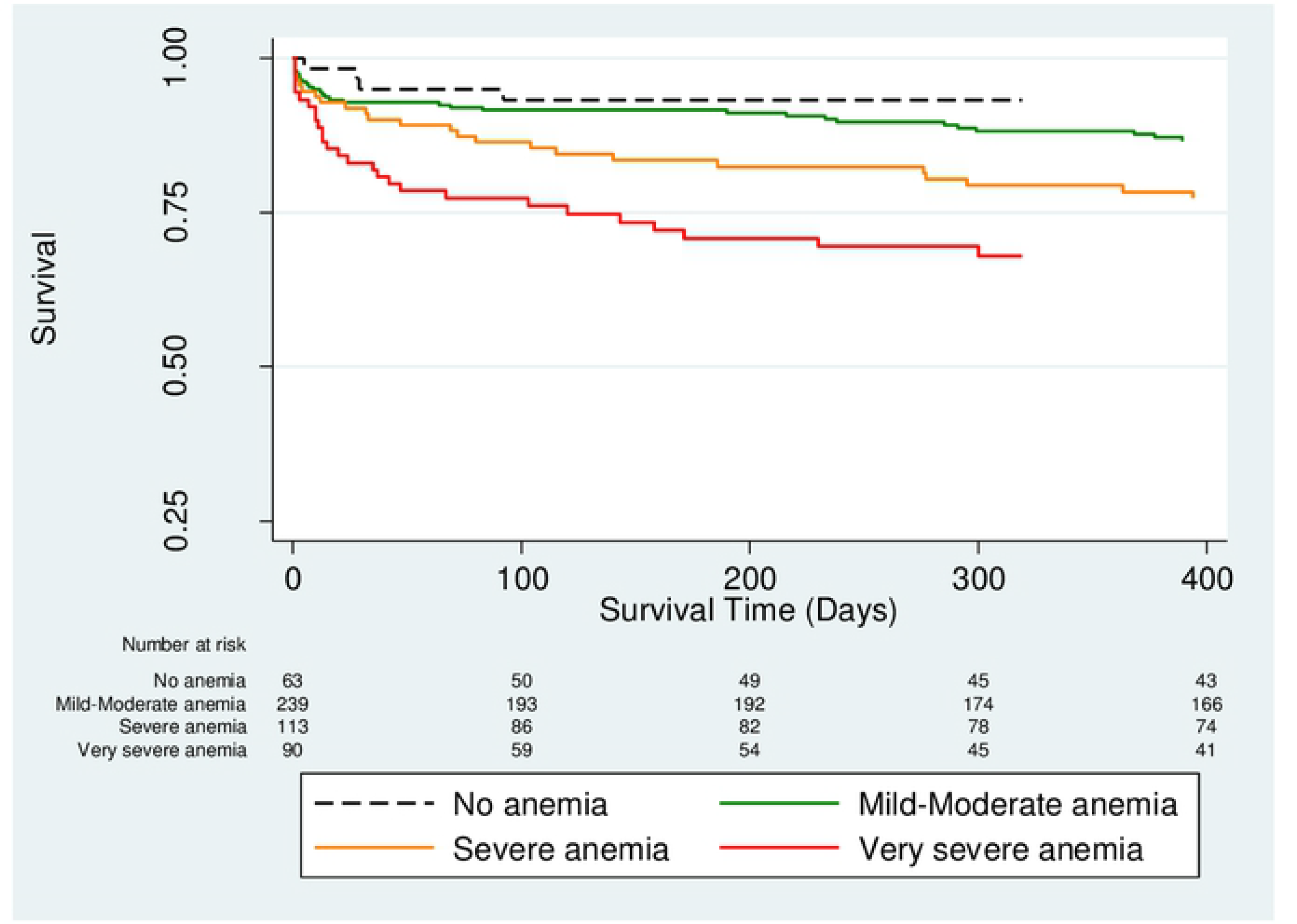
Kaplan Meier survival curves comparing severity of anemia (*p* < 0.001 by log-rank test).

### Factors associated with mortality

All variables listed in Table 1 were analyzed as possible predictors of mortality for children with very severe anemia (Tables 3 and 4). The significant independent predictors of mortality by multivariate Cox regression analysis were older age (HR 1.01, 95% Cl 1.00 – 1.03, *p* = 0.006), and reports of decreased urine output (HR 4.30, 95% Cl 1.04 – 17.7, *p* = 0.044). In regard to older age, overall mortality was significantly higher for children 5 years and older with very severe anemia compared to children under 5 years of age with very severe anemia, 17/37 (45.9%) vs. 10/53 (18.9%) respectively (HR 2.79, 95% Cl 1.27 – 6.10, *p* < 0.010).

**Table 3.**
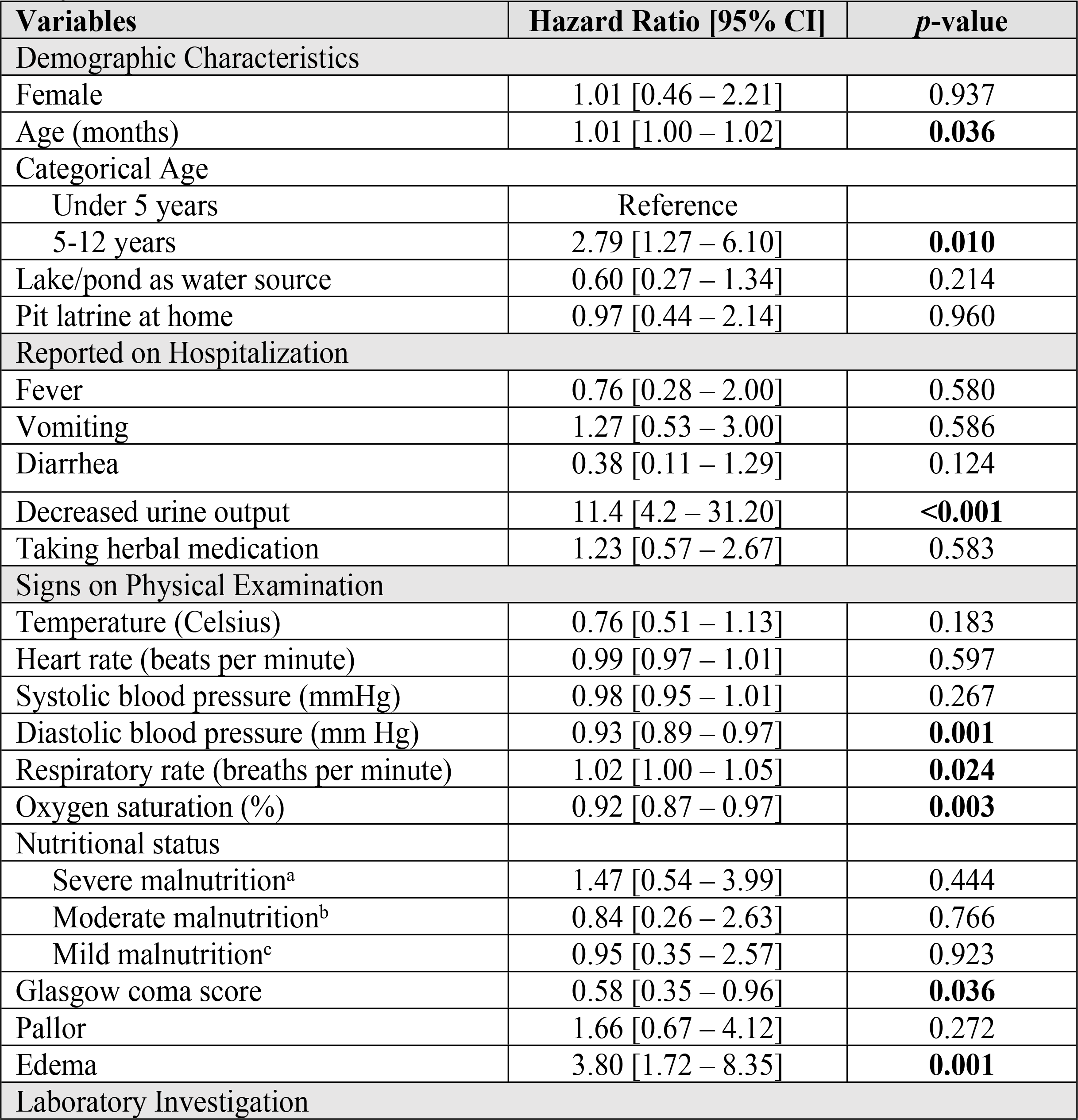
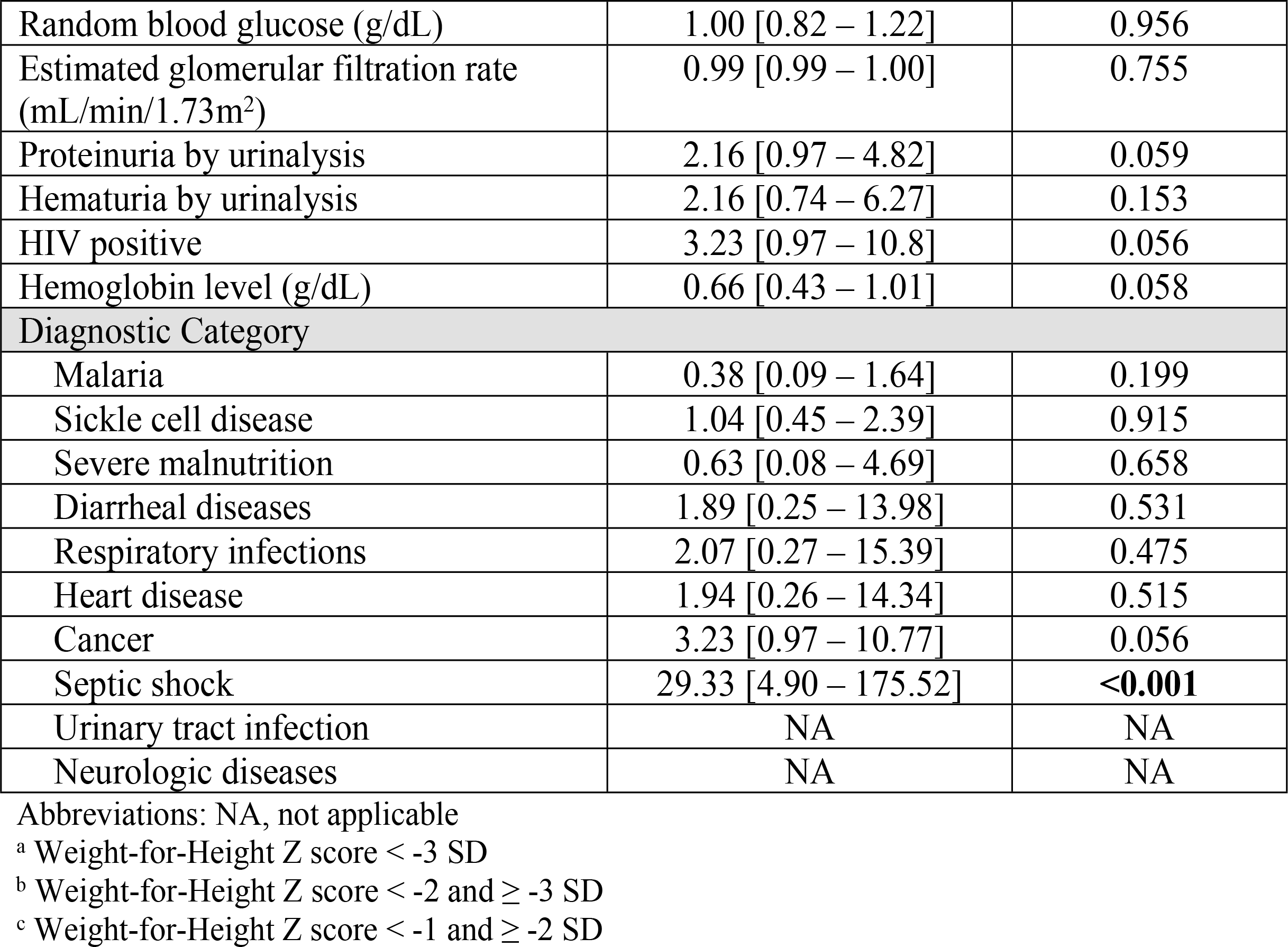
Predictors of mortality for children with very severe anemia by univariate analysis.

| Variables | Hazard Ratio [95% CI] | *p*-value |
| --- | --- | --- |
| Demographic Characteristics |  |  |
| Female | 1.01 [0.46 – 2.21] | 0.937 |
| Age (months) | 1.01 [1.00 – 1.02] | **0.036** |
| Categorical Age |  |  |
| Under 5 years | Reference |  |
| 5-12 years | 2.79 [1.27 – 6.10] | **0.010** |
| Lake/pond as water source | 0.60 [0.27 – 1.34] | 0.214 |
| Pit latrine at home | 0.97 [0.44 – 2.14] | 0.960 |
| Reported on Hospitalization |  |  |
| Fever | 0.76 [0.28 – 2.00] | 0.580 |
| Vomiting | 1.27 [0.53 – 3.00] | 0.586 |
| Diarrhea | 0.38 [0.11 – 1.29] | 0.124 |
| Decreased urine output | 11.4 [4.2 – 31.20] | **<0.001** |
| Taking herbal medication | 1.23 [0.57 – 2.67] | 0.583 |
| Signs on Physical Examination |  |  |
| Temperature (Celsius) | 0.76 [0.51 – 1.13] | 0.183 |
| Heart rate (beats per minute) | 0.99 [0.97 – 1.01] | 0.597 |
| Systolic blood pressure (mmHg) | 0.98 [0.95 – 1.01] | 0.267 |
| Diastolic blood pressure (mm Hg) | 0.93 [0.89 – 0.97] | **0.001** |
| Respiratory rate (breaths per minute) | 1.02 [1.00 – 1.05] | **0.024** |
| Oxygen saturation (%) | 0.92 [0.87 – 0.97] | **0.003** |
| Nutritional status |  |  |
| Severe malnutrition<sup>a</sup> | 1.47 [0.54 – 3.99] | 0.444 |
| Moderate malnutrition<sup>b</sup> | 0.84 [0.26 – 2.63] | 0.766 |
| Mild malnutrition<sup>c</sup> | 0.95 [0.35 – 2.57] | 0.923 |
| Glasgow coma score | 0.58 [0.35 – 0.96] | **0.036** |
| Pallor | 1.66 [0.67 – 4.12] | 0.272 |
| Edema | 3.80 [1.72 – 8.35] | **0.001** |
| Laboratory Investigation |  |  |
| Random blood glucose (g/dL) | 1.00 [0.82 – 1.22] | 0.956 |
| Estimated glomerular filtration rate (mL/min/1.73m<sup>2</sup>) | 0.99 [0.99 – 1.00] | 0.755 |
| Proteinuria by urinalysis | 2.16 [0.97 – 4.82] | 0.059 |
| Hematuria by urinalysis | 2.16 [0.74 – 6.27] | 0.153 |
| HIV positive | 3.23 [0.97 – 10.8] | 0.056 |
| Hemoglobin level (g/dL) | 0.66 [0.43 – 1.01] | 0.058 |
| **Diagnostic Category** |  |  |
| Malaria | 0.38 [0.09 – 1.64] | 0.199 |
| Sickle cell disease | 1.04 [0.45 – 2.39] | 0.915 |
| Severe malnutrition | 0.63 [0.08 – 4.69] | 0.658 |
| Diarrheal diseases | 1.89 [0.25 – 13.98] | 0.531 |
| Respiratory infections | 2.07 [0.27 – 15.39] | 0.475 |
| Heart disease | 1.94 [0.26 – 14.34] | 0.515 |
| Cancer | 3.23 [0.97 – 10.77] | 0.056 |
| Septic shock | 29.33 [4.90 – 175.52] | **<0.001** |
| Urinary tract infection | NA | NA |
| Neurologic diseases | NA | NA |

Abbreviations: NA, not applicable
<sup>a</sup> Weight-for-Height Z score < -3 SD
<sup>b</sup> Weight-for-Height Z score < -2 and ≥ -3 SD
<sup>c</sup> Weight-for-Height Z score < -1 and ≥ -2 SD

**Table 4.**
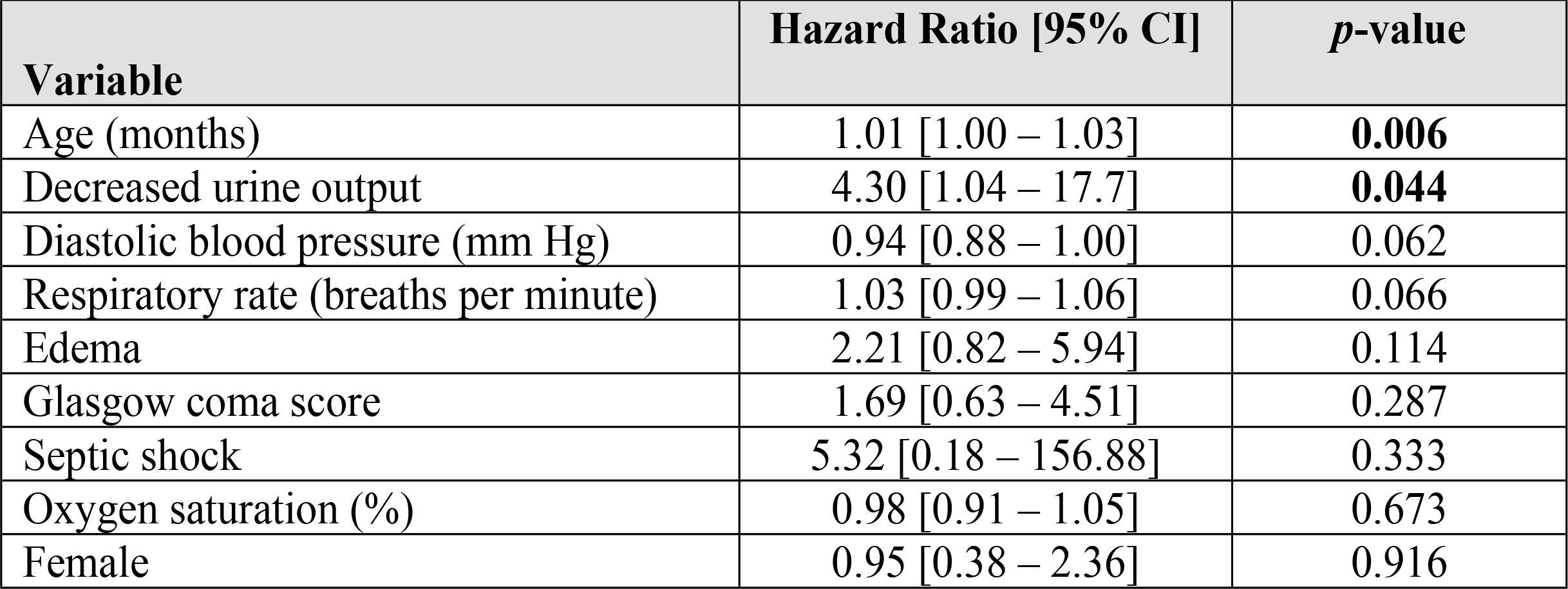
Predictors of mortality for children with very severe anemia by multivariate analysis.

Comparison of predictors for in-hospital and post-hospital mortality for children with very severe anemia are in S1 Table.

## Discussion

Children admitted to hospitals in Tanzania with very severe anemia suffer from strikingly high rates of mortality in the year after an index hospitalization. In our study, 30% of children with very severe anemia died within one year of hospitalization. Fifty-nine percent of those deaths occurred during the post-hospital period. This confirms and extends findings from a prior study of hospitalized Malawian children under the age of 5 years with hemoglobin less than 5.0 g/dL [9]. In this study, 17% of these children with hemoglobin less than 5.0 g/dL died in the 18 months after hospital admission. The majority these deaths (63%) occurred after hospital discharge. We demonstrate that even older children with very severe anemia suffer from alarmingly high rates of mortality in the post-hospital period. In fact, children 5 years of age and older with very severe anemia had even higher mortality than children under 5 years of age with very severe anemia. In regards to reducing child mortality, children under the age of 5 have received the vast majority of the attention. Efforts should be made to ensure older children also benefit from health policies and interventions [17–18].

For the deaths that occurred post-hospital for children with very severe anemia, more than half the deaths occurred within 2 months of discharge. This finding supports the idea of a period of vulnerability following discharge. A high post-hospital mortality rate in the early discharge period has also been shown for children with malaria, diarrhea and for general admissions [19–22]. This may be partly explained by the “post-hospital syndrome”, which has been described as an acquired transient period of vulnerability following discharge [23]. However, post-hospital mortality is not merely an issue of disease pathology, but also has a complex socioeconomic overlay in our setting. Problems related to poverty (e.g., funds for transportation to health facilities, paying for medical care), parental education, gender inequities (e.g., husbands making decisions while mothers having better understanding of the needs of their children), parental employment (e.g., work affecting ability to seek medical attention for their children) and seeking traditional medicine healers can reduce adherence to medications and clinic follow-up, and thereby lead to higher mortality [24–26]. We currently are piloting a post-hospital case management intervention at BMC addressing these socioeconomic factors to improve outcomes.

Low hemoglobin level is a strong predictor of mortality. In our study, children with very severe anemia were 4.3 times more likely to die, compared to children without any anemia. Our study also found for each 1 g/dL decrease in hemoglobin, the risk of death increased by 18%. This appears to be consistent with other published data. A meta-analysis of nearly 12,000 children from six African countries looking at the association between anemia and mortality reported an odds ratio of 0.76 (95% Cl 0.62 – 0.93), indicating that for each 1 g/dL decrease in hemoglobin, the risk of death increased by 24% [27]. For children with very severe anemia, attention on older aged children is critical, as increasing age appeals to be a predictor of mortality. One speculation is older children are more likely to have chronic diseases associated with anemia (e.g. sickle cell disease, cancer), compared to younger children who have not acquired chronic diseases yet. The ability for health facilities in low resource settings to diagnose children of all ages with anemia and treat with adequate blood supply if needed will be key to reducing child mortality. There is currently an ongoing multi-center trial in Uganda and Malawi that we hope will shed light into establishing best transfusion and treatment strategies in preventing mortality for children with anemia [28].

The prevalence of very severe anemia was found to be high among Tanzanian children who were hospitalized. In our study, nearly 20% of the children were found to have very severe anemia, and 40% of children had severe anemia or very severe anemia. The high prevalence of anemia appears to be consistent with a prior study done at our institution [29]. A possible explanation may be due to the high prevalence of malaria, sickle cell disease, soil transmitted helminths and nutritional deficiencies in our region [30–32]. By comparison to the general community or household level, the prevalence of severe anemia has been reported to be as high as 2.5% in East Africa and 3.4% in sub-Saharan Africa [33–34]. The high prevalence of anemia, whether for children hospitalized or in the community, highlights the continual need to make identification and treatment of anemia a public health priority.

Our study has limitations. We did not acquire a hemoglobin level at time of discharge, and there were limitations in our diagnostic facilities to determine causes of anemia at the time of our study (e.g., iron studies). We also had a higher lost to follow-up rate for children with very severe anemia. Children lost to follow-up may have been more likely to have died. This would make our findings underestimate the true risk that very severe anemia has on mortality.

In conclusion, we conducted a prospective cohort study of 505 children up to 12 years of age hospitalized with very severe anemia and followed for one year post-hospitalization in Tanzania. Nearly 30% of children hospitalized with very severe anemia died within one year. More than half of those deaths occurring after hospital discharge. Since anemia is extremely prevalent among both young and older African children, and the consequences of very severe anemia so deadly, prevention and treatment of anemia must be a high public health priority to reduce child mortality in Africa.

## Acknowledgements

The authors would like to thank the children and their caretakers involved in this study, the clinicians and nurses who cared for the children, and faculty in the Department of Pediatrics of BMC and STH.

## Supporting information

S1 Table. Predictors of in-hospital and post-hospital mortality of children with very severe anemia by univariate analysis.

